# Cholesterol-deficient T cell membrane-encapsulated STING agonists for tumor-targeted immunotherapy

**DOI:** 10.1101/2022.12.21.521349

**Authors:** Lin Li, Mengxing Zhang, Tiantian Liu, Zhirong Zhang, Zhenmi Liu, Ling Zhang

## Abstract

In recent years, STING agonists have shown promising results in enhancing tumor immunotherapies. Nanoparticle-based tumor targeting delivery of STING agonists is considered as an important way to improve the therapeutic efficacy and reduce the side effects of STING agonists. However, how to escape the clearance to nanoparticles by phagocytes in the blood while maintaining the tumor targeting efficiency of nanoparticles is still a big challenge. Herein, cholesterol-deficient membrane from bioengineered T cells overexpressing PD-1 encapsulated STING agonist SR-717 (a.k.a. COMs) was used to treat melanoma. Nanoparticles coated by these membranes displayed remarkably dropped clearance by monocytes in the blood in both animal and human blood comparing with nanoparticles coated by non-modified T cell membrane, while maintaining the high tumor cell targeting efficiency of COMs. In mice melanoma model, intravenous injected COMs successfully delivered SR-717 to tumor and activated STING pathways and the PD-1 on COMs blocked the up-regulated PD-L1 in tumor cells induced by SR-717. As a result, COMs stimulated strong tumor immune responses to inhibit melanoma recurrence when it combined with photothermal therapy (PTT). In summary, this study developed a highly effective bionic system that integrated STING activation and immunotherapy, and provided a simple and effective strategy to enhance performance of cell membrane-coated delivery systems in vivo.

## Introduction

It is known that immunotherapy benefit a lot for the treatment of tumors, and the inhibitors of PD-1 and PD-L1 have become the first-line standard treatment for patients with advanced tumors^1, 2^. However, most tumor patients could not benefit from immune checkpoint blockade (ICB) treatment as their tumors expressed low levels of PD-L1, and PD-L1 expression in the tumor immune microenvironment is recognized as both a prognostic and predictive biomarker in patients with melanoma^3-6^. Therefore, new combined immunotherapy strategies are urgently needed to improve effect of ICB^7^.

Recent studies shown that interferon gene (STING) plays a key role in initiating anti-tumor immunity^8^. The commonly used STING agonist cyclic guanosine monophosphate (cGAMP) could activate STING and induce the secretion of interferon (IFNs) to trigger anti-tumor immune responses, and the released IFNs was able to up-regulated PD-L1 in tumor cells, which can enhance the therapeutic effect of ICB^9-13^. However, cGAMP is unstable in vivo^14^, although the recently reported analog SR-717 has improved the stability to some extent^10^, it still lacks tumor targeting ability and may induce systemic toxicity. Thus, tumor targeting delivery of STING agonists is very necessary^15^.

Phenolic-metal nanoparticles are known to have excellent drug-trapping capacity but perform poorly tumor targeting in vivo^16^, here quercetin-ferrum nanoparticles (a.k.a. QFN) was utilized to delivery SR-717 (a.k.a. QFNs)^17^. As immunotherapy alone is difficult to cure the rapidly growing malignant tumors such as melanoma^18, 19^, the photothermal conversion effect of QFN may enhance the effect of immunotherapy against tumors in this study^17^. Cell membrane coated nanoparticles have been widely studied in recent years, and it has great advantages in targeted delivery of drugs^20^. In this study, we plan to use T cell membrane overexpressed with PD-1 to coat QFNs. We hope that PD-1 on T cell membrane can target and blockade PD-L1 on tumor cells, and thus combined STING agonist with ICB to treat tumors^21^. However, although nanoparticles coated with T cell membrane performed very well in tumor targeting in vitro, it was largely eliminated by monocytes in the blood, and good tumor targeting ability was not observed in this study.

When nanoparticles are intravenously injected into the blood, complement proteins deposit on the surface of nanoparticles in a process called opsonization^22, 23^. These proteins prime the particle for removal by immune cells like monocytes^24^. That’s maybe an important reason why some nanoparticles with targeting ligands that have shown strong tumor targeting ability in preclinical trials failed to show the corresponding ability in clinical study^24, 25^. Inhibiting the adsorption and activation of complement on the surface of nanoparticles is thought to be a promising strategy to increase the circulating time of nanoparticles in the blood, and scientists have made a lot of efforts to modify nanoparticles, such as polyethylene glycol (PEG) modification, sensitive structure modification or combined with complement inhibitors^23, 26-28^. However, how to selectively reduce^19^ the uptake of nanoparticles by phagocytes in the blood while maintain the tumor targeting ability of nanoparticles is still a big challenge, a simple and useful strategy is in an urgent need to enhance the tumor targeting of nanoparticles in vivo^29^.

The cell membrane used in delivery systems is mainly composed of phospholipid, cholesterol and proteins, but its morphology and composition are different from those of normal cells, and it is difficult for them to avoid being recognized as a foreign object and cleared by the phagocytes in the blood^30, 31^. It was reported that the elimination of liposomes from blood circulation was enhanced by increasing the cholesterol content in liposomes^32^. However, it still unknown that if reducing cholesterol content in cell membrane could improve the tumor targeting ability of cell membrane coated nanoparticles, and its mechanisms remain to be explored.

In this study, we found that T cell membrane with lower cholesterol content still maintains the targeting of CTTLL2-PD1 to tumor cells, and it can inhibit the uptake of nanoparticles by monocytes in the blood (∼50%). COMs performed better than common CTLL2-PD1 cell membrane or PEG-modified cell membrane coated nanoparticles in targeting tumors in vivo. COMs successfully delivered the STING agonist to tumor and activated STING pathways, while PD1 on the COMs surface reduced the up-regulated PD-L1 in tumor cells induced by STING agonist (Schem 1). Additionally, cholesterol-deficient B16F10 cell membrane-based nanoparticles also show improved tumor targeting ability in vivo, indicating that this strategy may also be suitable for other cell membrane coated nanoparticles.

## Results

### 1. Preparation and characterization of COMs

CTLL2 cells overexpressing PD-1 was constructed (CTLL2-PD1), and the expression of PD1 was significantly higher than that of CTLL2 cells (Figure 1A-B). Then, CTLL2-PD1 cells were treated with (2-hydroxypropyl)-β-cyclodextrin (β-CD), and then cell membrane was extracted for coating quercetin-ferrum nanoparticles (QFN), and QFN was used to capsulate SR-717 (QFNs) (Figure 1C). Results show that the content of cholesterol in cell membrane treated with β-CD (20 mM) is about 20% of the ordinary cell membrane (Figure 1D). TEM images show that obvious membrane like structure could be detected on the surface of CTLL2 cell membrane-coated QFNs (Ms), CTLL2-PD1 cell membrane-coated QFNs (OMs) and cholesterol-deficient CTLL2-PD1 cell membrane-coated QFNs (COMs) (Figure 1E). Fluorescence images of tumor cells show that the cell membrane of COMs had good co-localization with its core QFNs, indicating that the cell membrane was successfully coated in COMs (Figure 1F).

**Figure 1.**
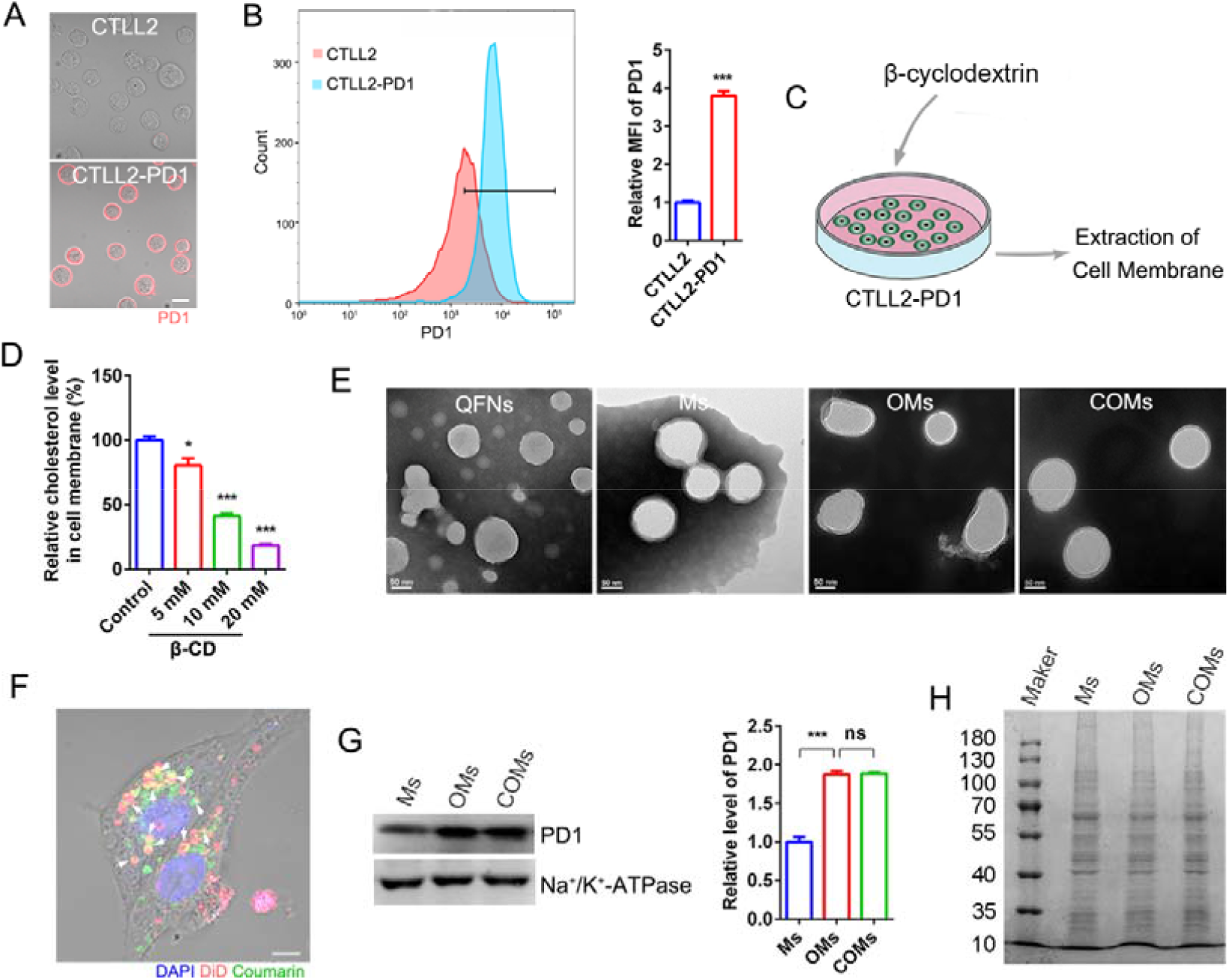
Preparation and characterization of COMs. (A) Representative fluorescence images of PD-1 on cell membrane. n=5. (B) Membrane PD-1 level detected by flow-cytometry, and the semi-quantitative analysis is shown on the right. n=3. (C) Schematic illustration of the way to prepare cell membrane with lower level of cholesterol. (D) Relative level of cholesterol in cell membrane after cells incubated with different concentration of (2-hydroxypropyl)-β-cyclodextrin for 0.5 h. (E) Representative TEM images of QFNs, Ms, OMs and COMs. (F) The co-localization of cell membrane (marked with DiD) and QFNs core (marked with coumarin) in COMs. B16F10 cells were incubated with COMs for 4 h. (G) WB analysis of PD1 and Na^+^/K^+^-ATPase in Ms, OMs and COMs. The semi-quantitative analysis is shown on the right. n=3. (H) Images of coomassie brilliant blue staining of SDS-PAGE gel.

The hydration particle sizes of Ms (T cells membrane encapsulated STING agonist SR-717), OMs (T cells membrane overexpressing PD-1 encapsulated STING agonist SR-717) and COMs were about 200 nm, and the zeta potential is about −10 mV (Figure S1, Figure S2). Western Blot (WB) results show that the expression of PD-1 in OMs and COMs was about twice as much as that of Ms, and no significant difference in the content of PD-1 in OMs and COMs was detected (Figure 1G). SDS-PAGE and proteomics results show that there was little difference in membrane protein expression between OMs and COMs (Figure 1H, Figure S3). In addition, photothermal conversion effect of QFN was utilized in this study, and we found that QFN, QFNs, Ms, OMs, COMs showed similar photothermal effect with 808 nm laser irradiation (Figure S4), which may enhance the immunotherapy against melanoma induced by COMs^17, 19, 33^.

### 2. COMs blocked the elevated PD-L1 induced by STING agonists in tumor cells

Compared with Ms and QFNs, COMs and OMs show stronger targeting to B16F10 cells, and there was no significant difference in these two groups in targeting tumor cells (Figure 2A). Fluorescence images also show similar results (Figure 2B). In addition, the uptake of COMs by B16F10^PD-L1 KO^ was significantly reduced, and the positive rate of tumor cells with particles was reduced from ∼45% to ∼25% (Figure S5A-B), indicating that the binding of PD-1 and PD-L1 played a key role in the tumor targeting of COMs.

**Figure 2.**
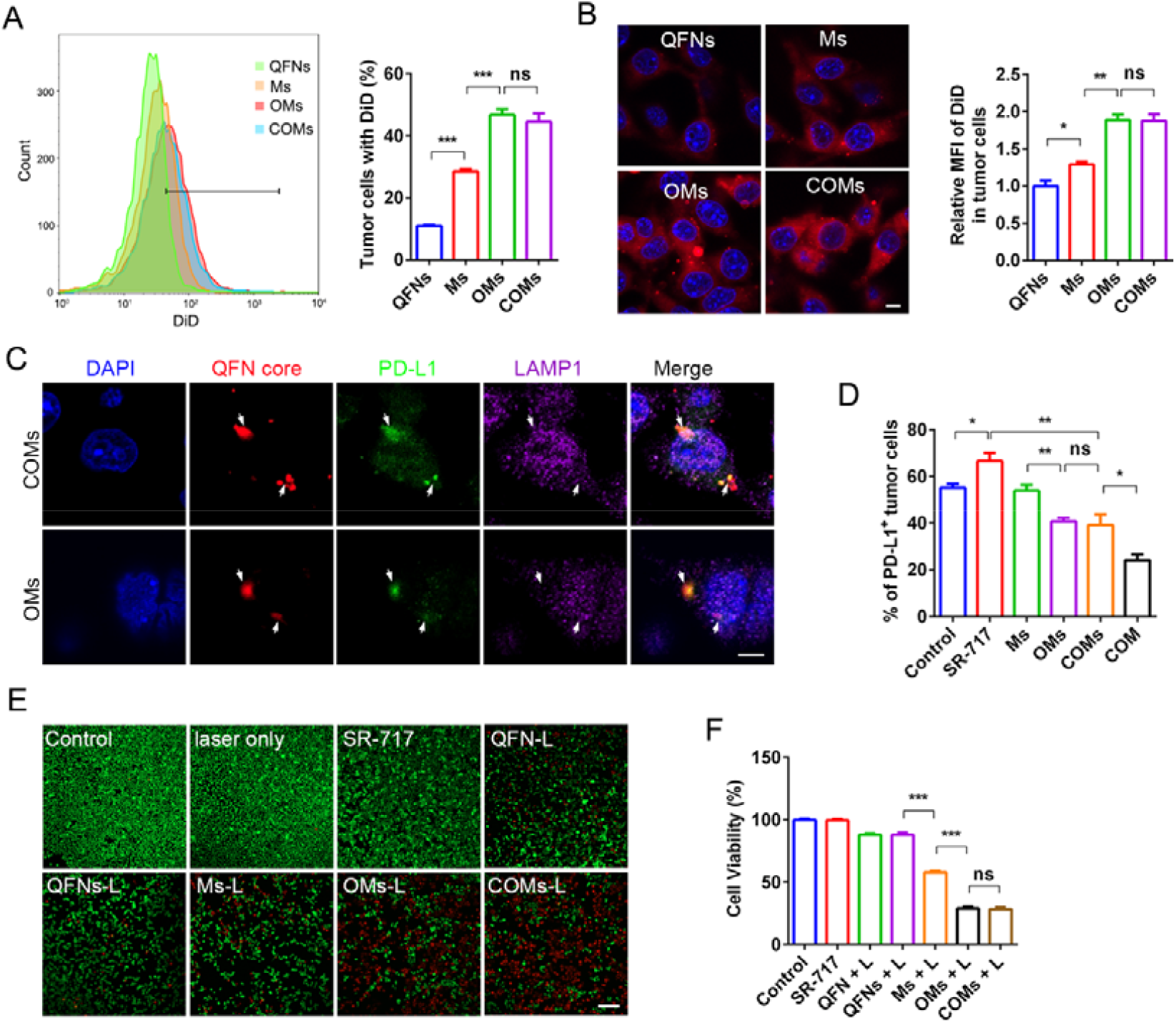
COMs blocked the elevated PD-L1 in tumor cells induced by STING agonists in vitro. (A) The uptake of QFNs, Ms, OMs and COMs by B16F10 cells detected by flow-cytometry, and the semi-quantitative analysis is shown on the right. Nanoparticles were marked with DiD. n=3. (B) Representative fluorescence images of B16F10 cells treated with QFNs, Ms, OMs and COMs. Nanoparticles were marked with DiD. The semi-quantitative analysis is shown on the right. n=3. (C) Representative images of the co-localization of QFN core, PD-L1 and LAMP1 in vitro. n=3. (D) Membrane PD-1 level detected by flow-cytometry. n=4. (E) B16F10 cells were treated with the indicated treatments, and then stained with fluorescein diacetate (green) and propidium iodide (red) and examined with fluorescence microscopy. n=5. (F) B16F10 cells were treated with the indicated treatments, and the cell viability was detected using CCK-8 kit.

In fact, both COMs and OMs could bind to PD-L1 in B16F10 cells and transport it to lysosomes for degradation (Figure 2C), which was consistent with previous report^21^. As mentioned, SR-717 can increase PD-L1 expression in tumor cells^10^, similar result was observed in this study (Figure 2D). Here, we found that OMs and COMs could reduce the expression of PD-L1 in tumor cells to a similar level (Figure 2D, Figure S6), indicating that COMs can blockade PD-L1 induced by SR-717 in tumor cells. Additionally, COMs and OMs show similar cytotoxicity to B16F10 cells under laser irradiation (Figure 2E, F).

Taken together, these results show that COMs could enhance the tumor targeting of nanoparticles by binding to PD-L1 in tumor cells, and PD-L1 is then degraded in lysosomes along with COMs, thereby reducing PD-L1 in tumor cells induced by SR-717. Moreover, COMs and OMs showed similar tumor targeting ability in vitro, indicating that reducing the cholesterol content of T cell membrane did not affect the tumor targeting of COMs in vitro.

### 3. COMs targeted to tumors in vivo

Encouraged by the good tumor targeting of COMs in vitro, we next study it in tumor bearing mice. Results show that the tumor targeting of COMs was significantly higher than that of QFNs, and fluorescence images of tumor sections also show similar results (Figure 3A-B, Figure S7). COMs could combine with PD-L1 in tumor cells in vivo and take it to lysosome for degradation, which was consistent with the results in vitro (Figure 2C, Figure 3C). COMs could also significantly increase the content of SR-717 in tumors (Figure 3D). However, OMs did not show only tumor targeting ability as it did in vitro, and there was no significant difference in the distribution of QFNs, Ms and OMs in tumors (Figure 3A-B). To improve the poor tumor targeting of OMs in vivo, we also tried to modify the cell membrane with PEG (PEG-OMs), hoping to improve the tumor targeting of OMs by prolonging the blood circulation time of nanoparticles^28, 34^. However, although PEG modified cell membrane could significantly reduce the uptake of nanoparticles by macrophages, it could also reduce the uptake of nanoparticles by B16F10 cells, and PEG-OMs failed to show tumor targeting ability in mice (Figure S8A-C).

**Figure 3.**
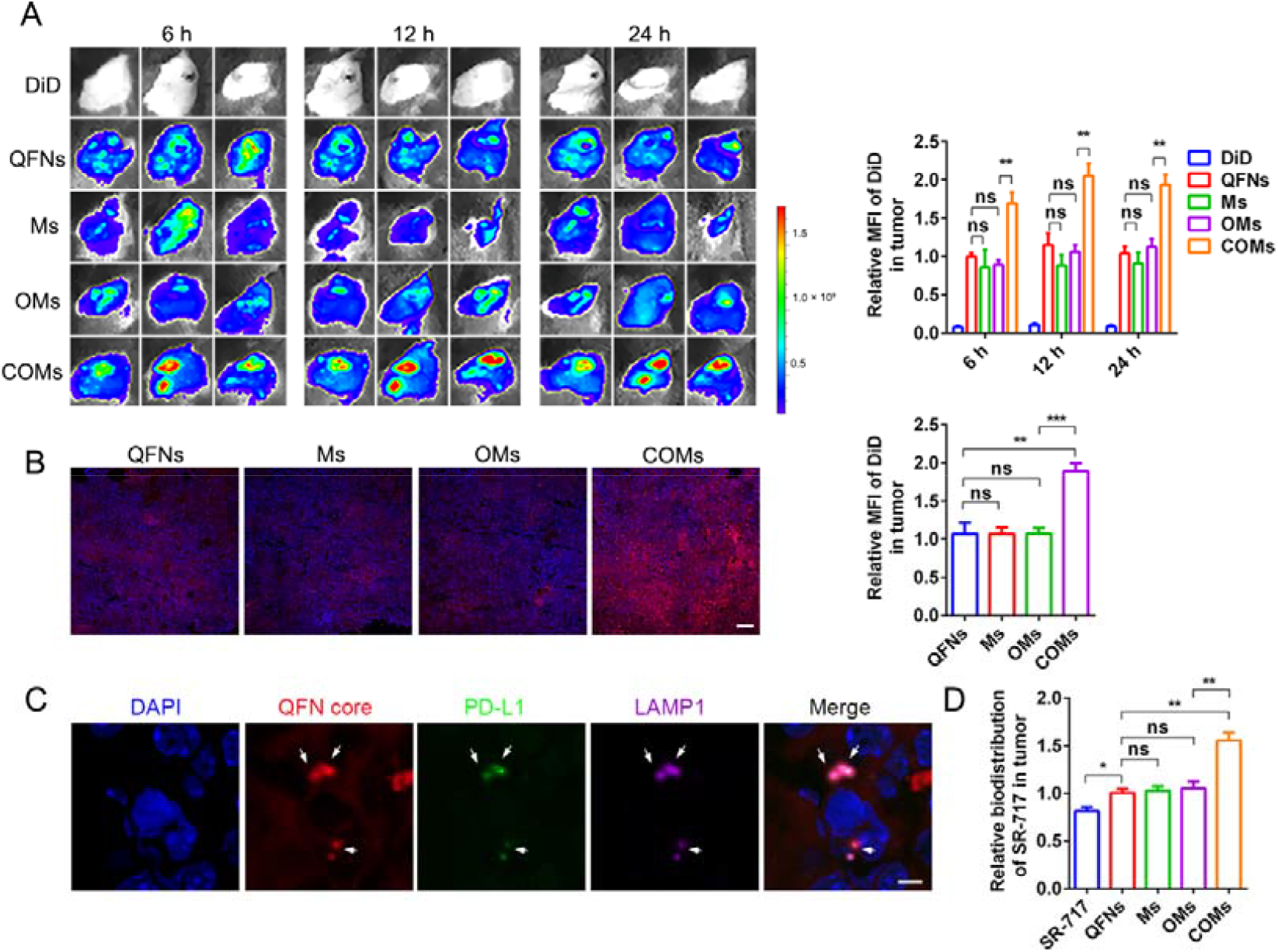
COMs could target to tumors in vivo. (A) Fluorescence images of tumor in mice treated with free DiD, QFNs, Ms, OMs and COMs(iv.). Nanoparticles were marked with DiD. The semi-quantitative analysis is shown on the right. n=3. (B) Representative images of the distribution of QFNs, Ms, OMs and COMs in tumors. The semi-quantitative analysis is shown on the right. n=3. (C) Representative images of the co-localization of QFN core, PD-L1 and LAMP1 in vivo. n=3. (D) Relative concentration of SR-717 in tumors measured by HPLC at 12 h post intravenously injecting of SR-717, QFNs, Ms, OMs and COMs. n=5.

In addition, CTLL2 cell membrane treated with β-CD could also improve the tumor targeting of QFNs (a.k.a. CMs), although it did not perform as good as that of COMs (Figure S9), which was consistent with the results in vitro study (Figure 2A), indicating that cholesterol-deficient membrane could show the tumor targeting ability of PD-1 ligands on nanoparticle in vivo. Besides, tumor cell membrane coated nanoparticles was able to target the same tumor^28^, although most of these works were did in node mice. We found that the B16F10 cell membrane treated with β-CD could also enhance the tumor targeting of B16F10 cell membrane coated nanoparticles to melanoma (Figure S10), indicating that this strategy is also applicable to other membrane coated nanoparticles.

### 4. The cholesterol-deficient cell membrane inhibited the clearance of COMS by monocytes in the blood

After intravenously injection, nanoparticles are circulated in the blood before they reach tumor site and captured by tumor cells. Considering that OMs and COMs have similar tumor targeting ability to tumor cells in vitro (Figure 2A-B), we mainly focus on the situation of OMs and COMs in the blood. Fluorescence images show that the tumor targeting ability of COMs was positively correlated with the concentration of β-CD, and the cell membrane with lower cholesterol content had stronger tumor targeting ability (Figure 1D, Figure 4A). The circulation time of COMs in blood was longer than that of OMs (Figure 4B). COMs with lower cholesterol content had higher blood retention in the blood at 4 h post injection (Figure 1D, Figure 4C). Phagocytes (mainly including monocytes, lymphocytes and neutrophils) in blood is an important player for the clearance of nanoparticles in the body^23, 26^. Then, we analyzed the uptake of COMs and OMs by phagocytes in the blood. Results show that the content of COMs in plasma was 1.5 times higher than that of OMs, and the content of COMs in monocytes was ∼50% of that of OMs (Figure 4E-G, Figure S11). The phagocytosis of COMs and OMs by neutrophils and lymphocytes was less than that of monocytes and no significant difference was observed in these two groups (Figure S12A-B), indicating that the clearance of OMs by monocytes in the blood might be an important reason for its failure in targeting tumors.

**Figure 4.**
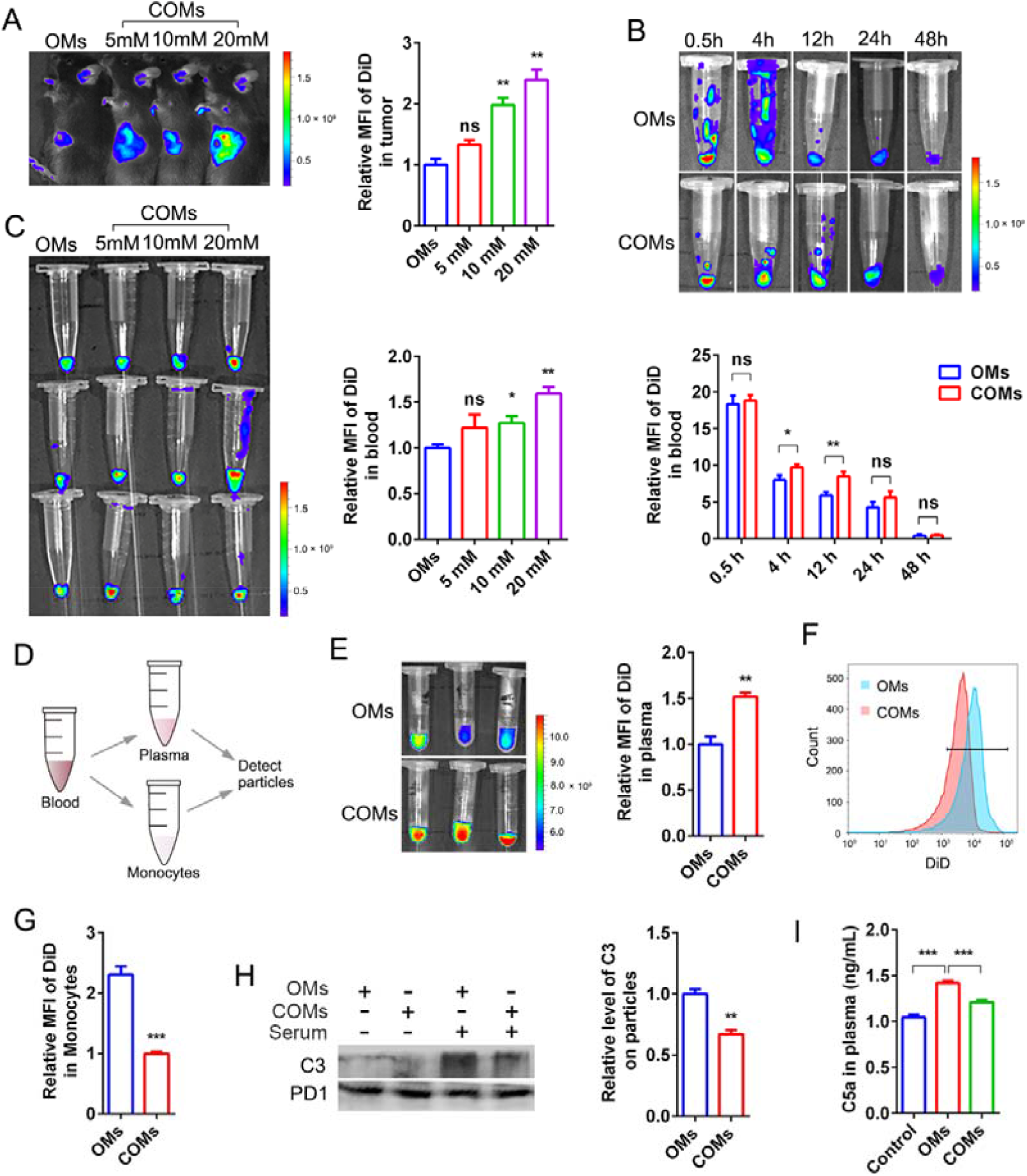
The cholesterol-deficient cell membrane inhibited the clearance of COMs by monocytes in the blood. (A) Fluorescence images of tumor in mice treated with COMs at 4 h post injection (iv.). The cell membrane used in COMs were obtained from CTLL2-PD1 cells incubated with different concentration of (2-hydroxypropyl)-β-cyclodextrin for 0.5 h. Nanoparticles were marked with DiD. The semi-quantitative analysis is shown on the right. n=3. (B) Representative fluorescence images of blood in tumor bearing mice. The semi-quantitative analysis is shown below. n=3. (C) The fluorescence images of blood in tumor bearing mice at 4 h post of injection (iv.) of COMs used in (A). The semi-quantitative analysis is shown right. n=3. (D) Schematic illustration of the experiment process. (E) Representative fluorescence images of plasma at 0.5 h post injection, and the semi-quantitative analysis is shown on the right. n=3. (F)(G) The uptake of OMs and COMs by monocytes in the blood at 0.5 h post injection, and the semi-quantitative analysis is shown in (G). n=3. (H) WB analysis of complement C3 in the serum bind to OMs and COMs, and the semi-quantitative analysis is shown on the right. n=3. (I) C5a in the plasma was measured using ELISA kit. n=3.

Complement proteins deposit on the surface of nanoparticles prime the particle for removal by immune cells like monocytes^22, 25, 26, 35^. We found that the adsorption of complement C3 on COMs in blood was significantly less than that of OMs, and the content of activated complement C5a in blood was also lower (Figure 4H-I), indicating that the weak adsorption and activation of complement system on COMs might be an important reason why COMs were less engulfed by monocytes in the blood^32^.

Then, the uptake of OMs and COMs by monocytes in human blood was examined. It is worth noting that after the nanoparticles incubated with human blood at 37 □ for 0.5 h, the uptake of COMs by monocytes in blood was also significantly lower than that of OMs, and more COMs were left in plasma (Figure S13), indicating that this strategy may have a good prospect of clinical transformation in the future.

### 5. COMs enhanced the anti-tumor immunity in mice

Encouraged by these results, we then tested the anti-tumor effect of COMs in melanoma in mice (Figure 5A). In order to enhance the therapeutic effect, photothermal therapy (PTT) is combined to enhance immunotherapy induced by COMs^17^. Results show that the tumor of mice treated with COMs was heated up rapidly with laser irradiation, and the tumor temperature was significantly higher than that of other groups (Figure S14). WB results show that COMs significantly increased the expression of p-TBK1 and p-IRF3 in tumors, indicating that COMs could activate STING signal pathway in tumors, while free SR-717, QFNs, Ms and OMs activated this pathway to a less extent (Figure 5B-D). Compared with SR-717, QFNs, Ms and OMs, COMs could also significantly reduce the expression of PD-L1 in tumor cells (Figure 5E,Figure S15). After photothermal/immunotherapy, COMs significantly increased the content of CD4^+^T cells, CD8^+^T cells, and M1 macrophages in tumors, and could also promote the maturation of DC cells in the tumor draining lymph nodes (Figure 5F-I, Figure S16). The immunofluorescence staining images show that COMs could significantly increase IFN-γ in tumors (Figure 5J). Not surprisingly, ∼80% of the mice in COMs group were cured after COMs-based photothermal/immunotherapy (Figure 5K-L). The effects of QFNs, Ms and OMs were similar, and 30% of tumors had no recurrence. Compared with QFN group, the tumor recurrence rate in QFNs group was improved, indicating that SR-717 could promote tumor therapy by activating tumor immune microenvironment (Figure 5K-L). To sum up, COMs could deliver SR-717 to tumors and block PD-L1 of tumor cells, which enhanced the anti-tumor immunity in mice.

**Figure 5.**
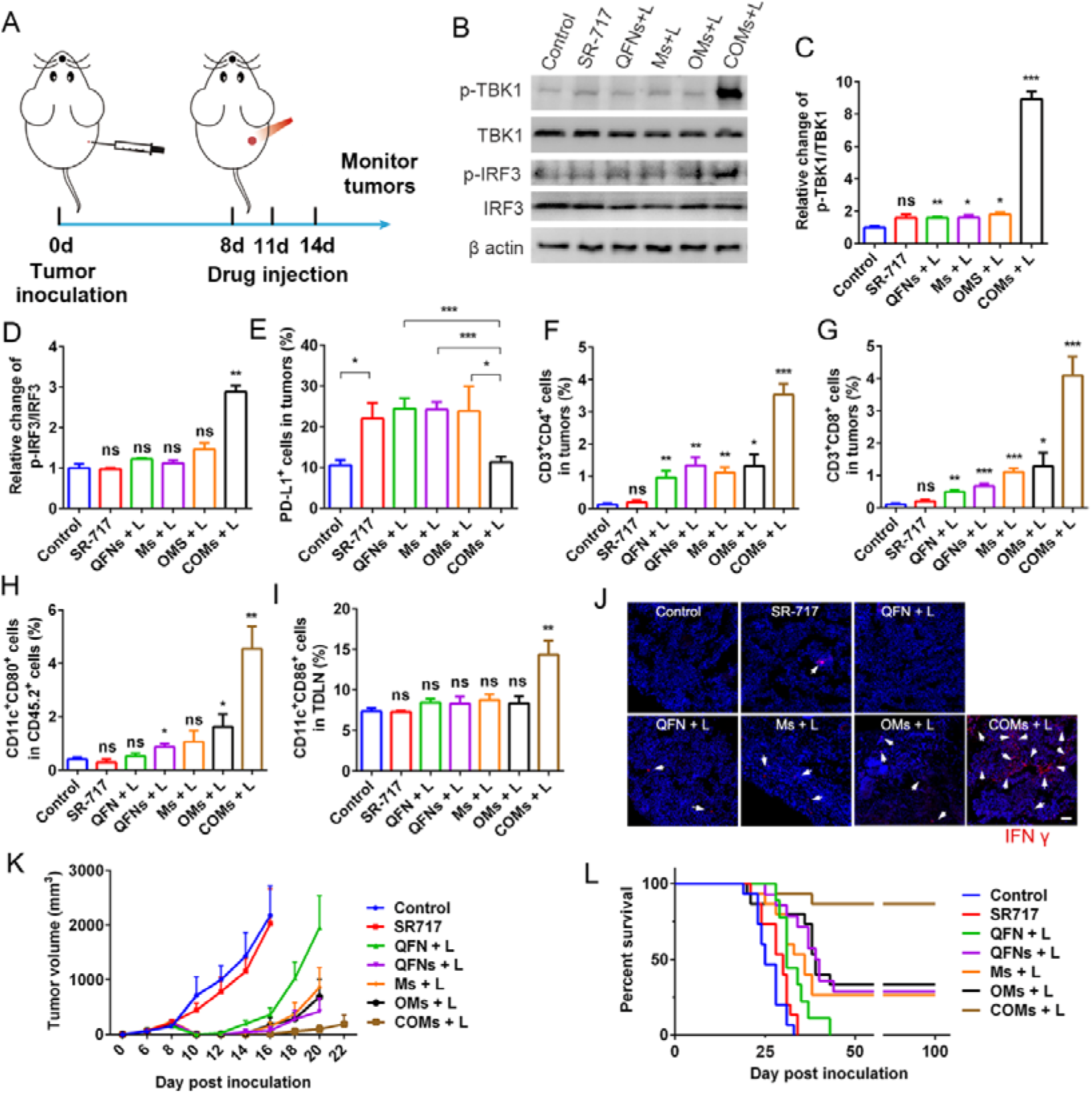
COMs enhanced the anti-tumor immunity in mice. (A) Schematic illustration of the experiment process. (B)(C)(D) WB analysis of p-TBK1, TBK1, p-IRF3 and IRF3 in tumors (B), and the semi-quantitative analysis is shown in (C)(D). n=3. (E) Flow-cytometric analysis of PD-L1 in tumors at 7 days post laser irradiation. n=5. (F) Flow-cytometric analysis of CD3^+^CD4^+^ in tumors at 7 days post laser irradiation. n=5. (G) Flow-cytometric analysis of CD3^+^CD8^+^ in tumors at 7 days post laser irradiation. n=5. (H) Flow-cytometric analysis of CD11c^+^CD80^+^ in CD45.2^+^ cells in tumors at 7 days post laser irradiation. n=5. (I) Flow-cytometric analysis of CD11c^+^CD86^+^ in tumor-draining lymph nodes. n=5. (J) Representative fluorescence images of IFN γ in tumors at 7 days post laser irradiation. n=5. (K)(L) Tumor growth (K) and survival curve of mice after different treatments (L). n=12.

Additionally, the biocompatibility of COMs was evaluated^17, 36^. Body weight of mice was unaffected by COMs (Figure S16). Hematoxylin-eosin staining of organs show that COMs did not induce clear kidney injury, pulmonary toxicity, cardiac injury or inflammatory infiltrates in the spleen (Figure S17A). The number of red blood cells, platelets and white blood cells in the COMs-treated group was similar to that of the control group (Figure S17B). These results indicate that COMs are relative safe in mice.

## Discussions and conclusions

In this study, QFN was utilized to wrap SR-717, and cholesterol-deficient T cell membrane was utilized to coat QFNs to promote the tumor targeting delivery of SR-717. Immunotherapy along is difficult to cure melanoma as it grows rapidly and has high degree of malignancy^5, 6^. Thus, the photothermal conversion effect of QFN was used to enhance the effect of immunotherapy against melanoma in this study^17^.

It is difficult for people to find a balance between prolonging the circulating time of nanoparticles and retaining the tumor targeting of nanoparticles^28^ (Figure S8). Many nanoparticles that showed targeted effect in preclinical studies failed to show the corresponding ability in clinical trials, and the clearance of nanoparticles by phagocytes in blood may be an important reason^22, 24, 37, 38^. In fact, the shape and size of the cell membrane wrapping nanoparticles is different from that of the normal cells in the blood, and it is difficult for them to avoid being recognized as a foreign object and cleared by phagocytes in the blood^30, 31^. Although CTLL2 cell membrane coated nanoparticles showed tumor targeting in the BALB/c mince bearing 4T1 tumors^21^, it may be owing to the clearance of nanoparticles by monocytes in the blood of BALB/c mice and C57 mice is different. Considering that human body has a complex immune system, the cell membrane coated nanoparticles may also could not exhibit good targeting ability in future clinical study^25^. This study found that lowering the cholesterol of the cell membrane could inhibit the clearance of nanoparticles by monocytes in human blood (Figure S13), thus providing a simple strategy to improve the targeting of cell membrane coated nanoparticles in clinical study.

The highest concentration of (2-hydroxypropyl)-β-cyclodextrin (β-CD) used in this study was 20 mM, and we did not further increase its concentration because higher concentration of β-CD may affect the cell activity of CTLL2-PD1 and affect the targeting ability of COMs. In addition, as cholesterol is an important for the stability of cell membrane, to leave a little content of cholesterol in cell membrane may help to maintain the morphology of COMs. Additionally, the detailed mechanism of COMs to adsorb less complement needs to be studied in the future^25, 32^.

In addition to treating tumors, this strategy may also be applicable to other cell membrane to treat other diseases (Figure S10), such as improving the targeting of neutrophil membrane-coated nanoparticles to arthritis^39^, or increasing the retention time of some nanoparticles that need to circulate in the blood for a long time. The cholesterol deficient cell membrane may also be used to coat bacteria^40^, reduce the recognition and elimination of bacteria by phagocytes in the blood, and enhance the enrichment of bacteria in target sites (such as tumors)^41^.

In summary, STING agonists have shown good therapeutic effects in many preclinical studies of tumors. However, there is still a lack of a simple and efficient tumor targeting delivery system for STING agonists to enhance its therapeutic effect. In this study, SR-717 was coated with cholesterol deficient T cell membrane, which reduced the adsorption and activation complements on COMs while retaining its tumor targeting ability, and reduce the clearance of COMs by monocytes in the blood. Thus, COMs showed good tumor targeting in mice, successfully delivered SR-717 to tumors, and activated the tumor STING signal pathway. Moreover, PD-1 overexpressed in COMs blockaded PD-L1 in tumor cells induced by SR-717, which enhanced the infiltration of immune cells in tumors. Finally, 80% of tumor bearing mice treated with COMs were cured with the combined PTT. This study provides an effective tumor targeting delivery system for SR-717, and this study has important implications for prolonging the circulation time in blood and enhancing the targeting ability of nanoparticles.

## Materials and methods

### Reagents

Quercetin, FeCl_2_.4H_2_O, (2-hydroxypropyl)-β-cyclodextrin, Coumarin, DiD dyes, EDTA and Nafamostat mesylate were brought from MACKLIN in China; Protease Inhibitor Cocktail, SR-717 and DAPI were purchased from MedChemExpress; PLGA40K-COOH(50:50) was obtained from Chongqing Yusi Pharmaceutical Technology in China; antibodies for flow cytometry assay were purchased from Biolegend; anti-PD1, anti-PD-L1, anti-C3, anti-TBK1 and anti-β actin for western blot were obtained from Abcam; anti-p-TBK1 and anti-p-IRF3 were purchased from CellSignalingTechnology; anti-IRF3 was purchased from Abclonal Technology in China; anti-Na^+^/K^+^ ATPase was purchased from HuaBio in China; anti-LAMP1 was purchased from Proteintech in China; RPMI-1640 medium and trypsin were purchased from KeyGEN BioTECH in China; Mouse Monocyte Extraction Kit, Human Monocyte Extraction Kit, Mouse Neutrophil extraction kit, Mouse Lymphocyte extraction kit and Puromycin was purchased from Solarbio in China; Cholesterol Assay Kit was obtained from ApplyGEN in China; C5a ELISA Kit was obtained from Redotbiotech; DSPE-PEG2000 was purchased from Ponsure Biological.

### Animals and cells

Male C57BL/6 mice (6–8 weeks) were obtained from GemPharmatech in China. All animal experiments were approved by the Institutional Animal Care and Ethics Committee of Sichuan University. B16F10 was brought from American Type Culture Collection and were cultured in complete 1640 medium. B16F10^PD-L1 KO^ was established in our previous work^17^, and it was cultured in complete 1640 medium; CTLL2 was purchased from Procell Life Science&Tech-nology.Co.,Ltd, and it was cultured in complete 1640 medium with IL-2 (100 U ml^−1^); CTLL2-PD1 was established by Shanghai Genechem in China, and it was cultured in complete 1640 medium with IL-2 (100 U ml^−1^). All cells were maintained in a humidified atmosphere incubator containing 5% CO_2_ at 37 °C.

### Preparation of QFNs

Oil phase: Quercetin (1 mg), FeCl_2_.4H_2_O (1 mg), SR-717 (1 mg) and PLGA (5 mg) were dissolved in ethanol (50 μL) and CH_2_Cl_2_ (1 mL), stirred for 5 min. Water phase: Polyvinyl alcohol (PVA, 20 mg) and tween 80 (10 mg) were dissolved in water (2 mL). Mix oil phase and water phase, treated with probe ultrasonic for 10 min (300 W), then remove dichloromethane with rotary evaporator. To prepare DiD-labeled nanoparticles, DiD (60 μg) was dissolved in the oil phase.

### Reduce the cholesterol level in cells

CTLL2-PD1 cells cultured in the complete 1640 medium with IL-2 were washed with PBS, and then cells were incubated with different concentration of (2-hydroxypropyl)-β-cyclodextrin in 1640 medium (without serum) for 0.5 h, 20 mM of (2-hydroxypropyl)-β-cyclodextrin was used in most experiments otherwise stated. The cell membrane was then prepared to make COMs.

### Derivation of cell membrane

To prepare cell membrane, previously reported methods were used with some modification^31, 42^. Briefly, CTLL2, CTLL2-PD1 and CTLL2-PD1 treated with β-CD were collected and disrupted in hypotonic lysing buffer (pH 7.4, 20 mM Tris-HCl, 10 mM KCl, 2 mM MgCl_2_, Protease Inhibitor Cocktail) at 4 □ for 30 min. Destroyed cells in ice bath using a probe ultrasonic instrument (150 W, 3 min). Then centrifuged at 2×10^4^ g for 20 minutes to remove organelles and large particles. The supernatant was centrifuged at 1.0×10^5^ g for 40 min to harvest cell membrane. The pellet was then washed once in 10 mM Tris-HCl pH = 7.5 and 1 mM EDTA. The content of cholesterol in cell membrane was assayed using Cholesterol Assay Kit (ApplyGEN). *Preparation and characterization of Ms, OMs and COMs* QFNs (1 mL) was mixed with each cell membrane (derived from 4×10^7^ cells), and sonicated using a probe ultrasonic instrument (100 W, 1 min) in ice bath, then extruded through 400 nm and 200 nm polycarbonate membranes with extruder for 20 passes. Size of Ms, OMs and COMs were examined using Malvern Zetasizer ZS90 and TEM.

### Preparation of PEG-OMs

To prepare PEG-OMs, previously reported method was used with some modification^28^. Briefly, membrane (derived from 4×10^7^ CTLL2-PD1 cells), 180 μg of DSPE-PEG2000 were mixed and were physically extruded through a 200 nm polycarbonate membrane for seven passes. The membrane vesicles were mixed with QFNs and sonicated using a probe ultrasonic instrument (100 W, 1 min) in ice bath, then extruded through 400 nm and 200 nm polycarbonate membranes with extruder for 20 passes.

### Immunofluorescence staining

To stain PD-1 on cell membrane, cells were washed with cold PBS, and then incubated with anti-PD-1 at 4 °C for 45 min. Cells were then observed with confocal laser scanning microscope.

To stain IFN-γ, PD-L1, LAMP1 in melanoma, tumors were fixed with 4% paraformaldehyde for 72 h. Tumors were gradually dehydrated in 15% and 30% sucrose solution, and then frozen sections (10 μm) were prepared. Then the sections were permeabilized with 0.1 % Triton X-100 and incubated with anti-IFNγ (APC), anti-PD-L1 (FITC) at 4°C for 45 min. To stain LAMP1 in sections, sections treated with Triton X-100 were incubated with anti-LAMP1 for 1.5 h at room temperature, then washed with PBS for 3 times. PE-conjugated secondary antibody was then added and incubated for 1.5 h at room temperature, then washed with PBS for 3 times. Images were taken using confocal laser scanning microscope.

### Uptake of QFNs, Ms, OMs and COMs by B16F10 cells in vitro

B16F10 cells (2×10^5^) were inoculated cells on 12 well plate. 24 h later, cells were treated with QFNs, Ms, OMs and COMs (marked by DiD) for 2 h. Cells were collected and the uptake of nanoparticles by cells was detected with flow cytometry and confocal laser scanning microscope.

### Biodistribution of DiD, QFNs, Ms, OMs and COMs in mice bearing melanoma

Subcutaneously inoculated B16F10 cells (0.7×10^6^ cells/mouse) in male C57BL/6 mice. At 10 days post cell inoculation, mice were intravenously injected with DiD, QFNs, Ms, OMs and COMs (marked by DiD, 6 μg/mice). Fluorescence images of tumors in mice were taken at 6 h, 12 h and 24 h post injection.

### The uptake of OMs and COMs by phagocytes in the blood, and C5a in the blood

Subcutaneously inoculated B16F10 cells (0.7×10^6^ cells/mouse) in male C57BL/6 mice. At 10 days post cell inoculation, mice were intravenously injected with OMs and COMs (marked by DiD, 6 μg/mice). Mice were sacrificed (0.5 h post injection) and the blood were collected using anticoagulant tube containing EDTA and nafamostat mesylate. Monocyte, neutrophil and lymphocyte in the blood were separated using Mouse Monocyte Extraction Kit, Mouse Neutrophil extraction kit and Mouse Lymphocyte extraction kit. Then, the uptake of OMs and COMs by monocytes was measured by flow cytometry. The plasma was collected and the fluorescence images were taken. The C5a in the plasma was examined using ELISA Kit.

To study the uptake of OMs and COMs in human blood, blood was obtained from healthy human subjects (according to local approved protocols and with individual consent) into blood tubes containing the anticoagulant lepirudin, which does not affect the complement system^22^. Then, 0.5 mL of fresh blood and 50 μL of DiD-labeled OMs and COMs were mixed and incubated in 37 [ water bath in dark for 0.5 h. Monocytes in blood were then separated using Human Monocyte Extraction Kit, and the uptake of nanoparticles by monocytes was measured with flow cytometry. The plasma was collected and the fluorescence images were taken.

### The bonding of complement C3 to nanoparticles

The serum of mice was collected, and 0.3 mL of serum was mixed with 0.1 mL of OMs or COMs, and incubated in 37 □ for 0.5 h. Then centrifuged at 2×10^4^ g for 30 min to separate OMs and COMs, washed with cold PBS. WB experiment was used to assay the bonding of complement C3 to nanoparticles.

### In vivo anticancer treatment

Subcutaneously inoculated B16F10 cells (0.7×10^6^ cells/mouse) in male C57BL/6 mice (n=12). On the 8th, 11th and 14th days after tumor cells were inoculated, drugs were intravenously injected (SR-717 10 mg/kg). The tumor was irradiated with 808nm laser for 5 minutes (1.0 W/cm^2^) at 12 h post the first injection.

### Statistical analysis

Statistical analysis was performed using GraphPad prism software. An unpaired two-tailed t-test was used to compare between two groups. All data are presented as mean ± s.e.m. Differences are considered statistically significant with *p < 0.05, **p < 0.01 and ***p < 0.001, ns, no significant difference.

## Acknowledgments

We acknowledge the support from the National Science Fund for Excellent Young Scholars (No. 82022070) and the Regional Innovation and Development Joint Fund (No. U20A20411)

## Author contributions

Lin Li designed this research. Lin Li and Mengxing Zhang performed and analyzed most experiments. Zhirong Zhang, LeYao Fu, Jing Li, Tiantian Liu, Zhenmi Liu and Ling Zhang assisted in analyzing the results. All authors have given approval to the final version of the manuscript.

## Figures

**Schem 1.**
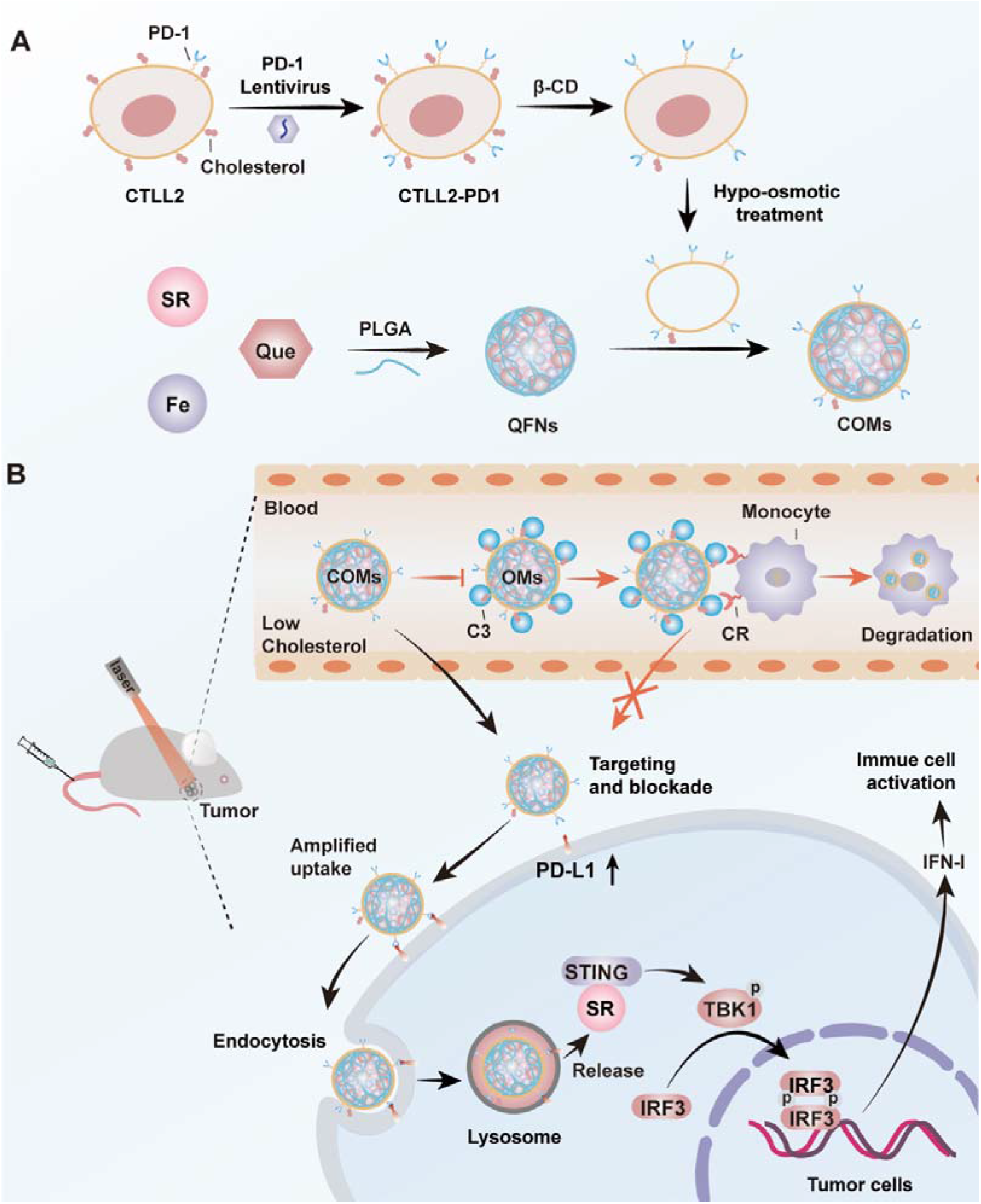
(A) Schematic illustration of the preparation of COMs. (B) COMs with low level of cholesterol enhanced the tumor-targeted delivery of SR-717.

## Supporting Information

**Figure S1.**
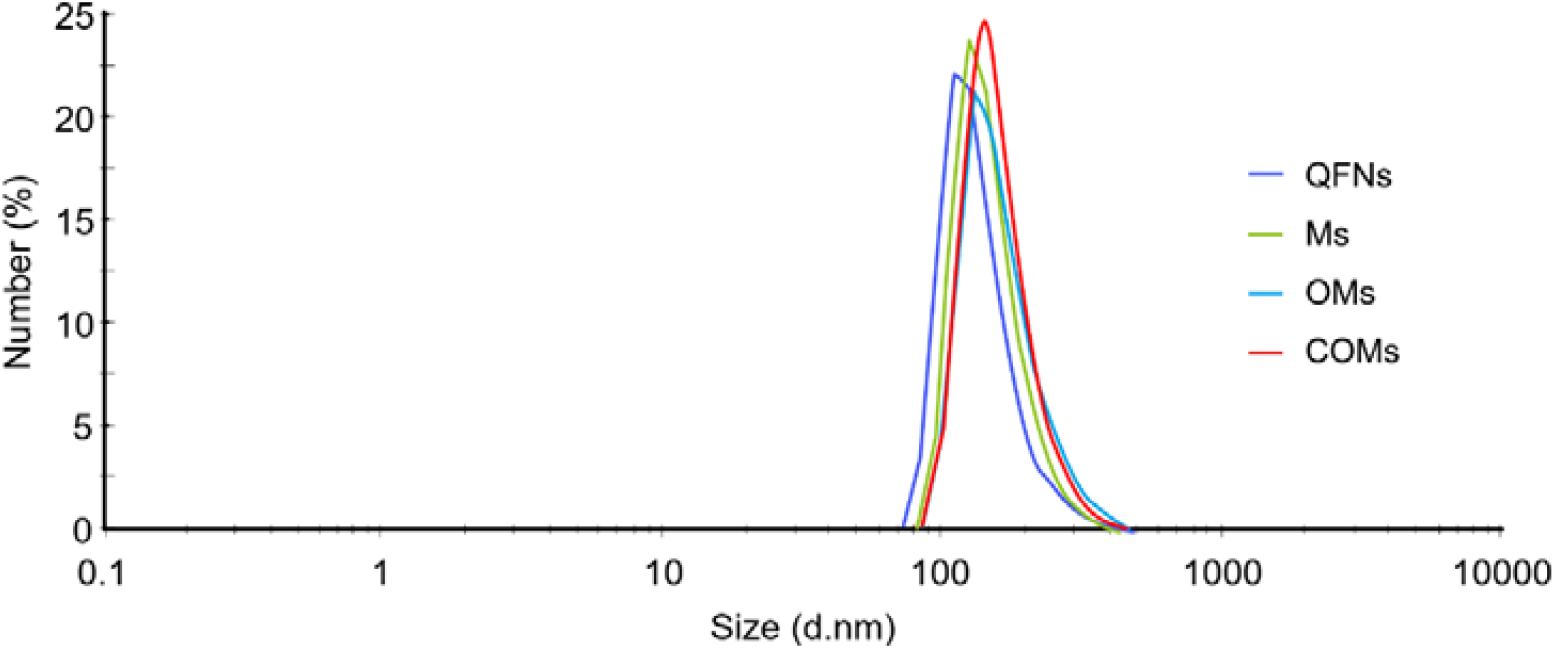
Sizes of QFNs, Ms, OMs and COMs.

**Figure S2.**
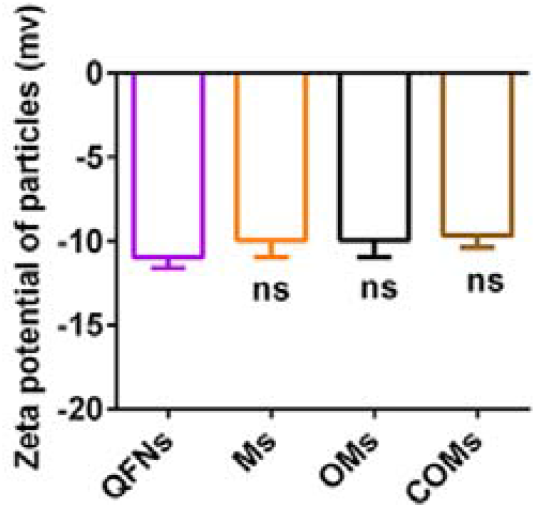
Zeta potential of QFNs, Ms, OMs and COMs. n=3.

**Figure S3.**
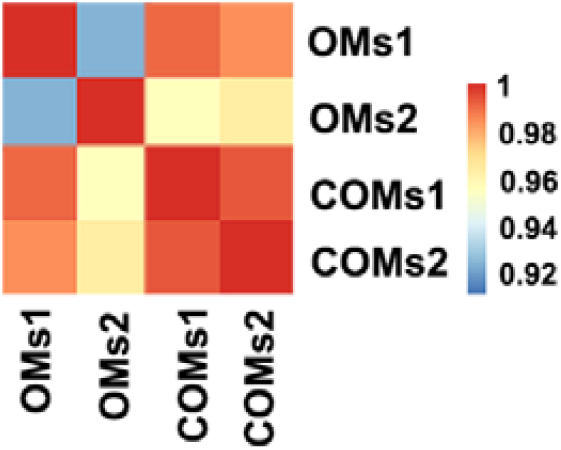
Protein expression difference in OMs and COMs.

**Figure S4.**
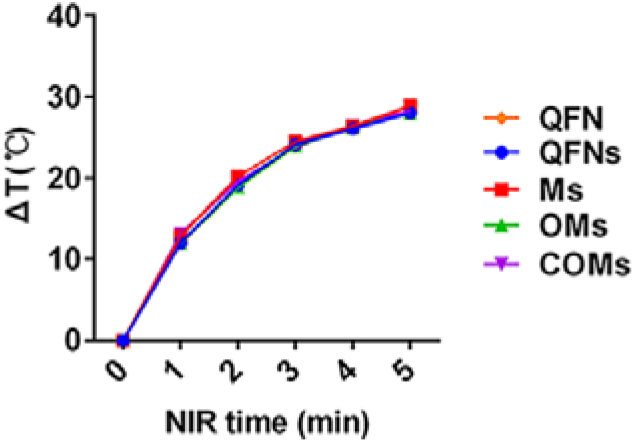
Temperature curves of QFN, QFNs, Ms, OMs and COMs with laser irradiation (1.0 W/cm^2^). The concentration of QFN is 200 μg/mL in all groups. n=3.

**Figure S5.**
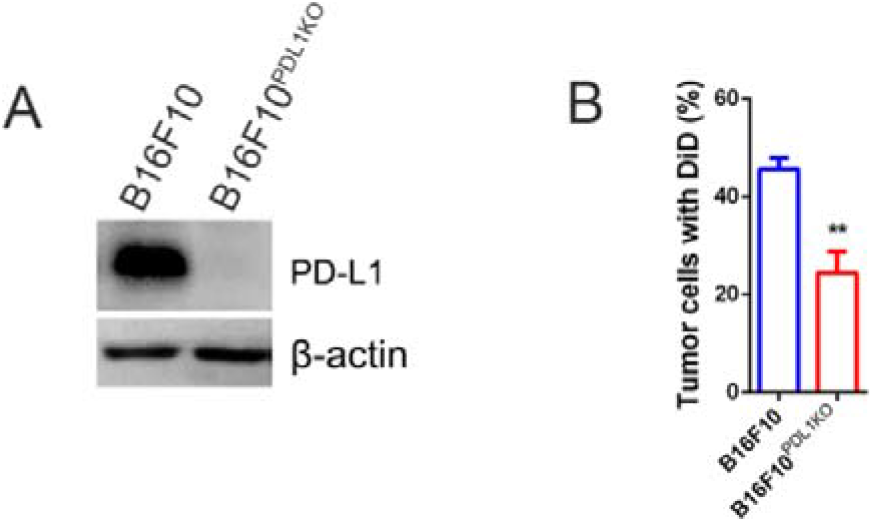
The uptake of COMs by B16F10^PDL1 KO^ cells. (A) WB analysis of PD-L1 in B16F10 cells and B16F10^PDL1 KO^ cells. (B) The uptake of COMs by B16F10^PDL1 KO^ cells was detected by flow-cytometry. n=5.

**Figure S6.**
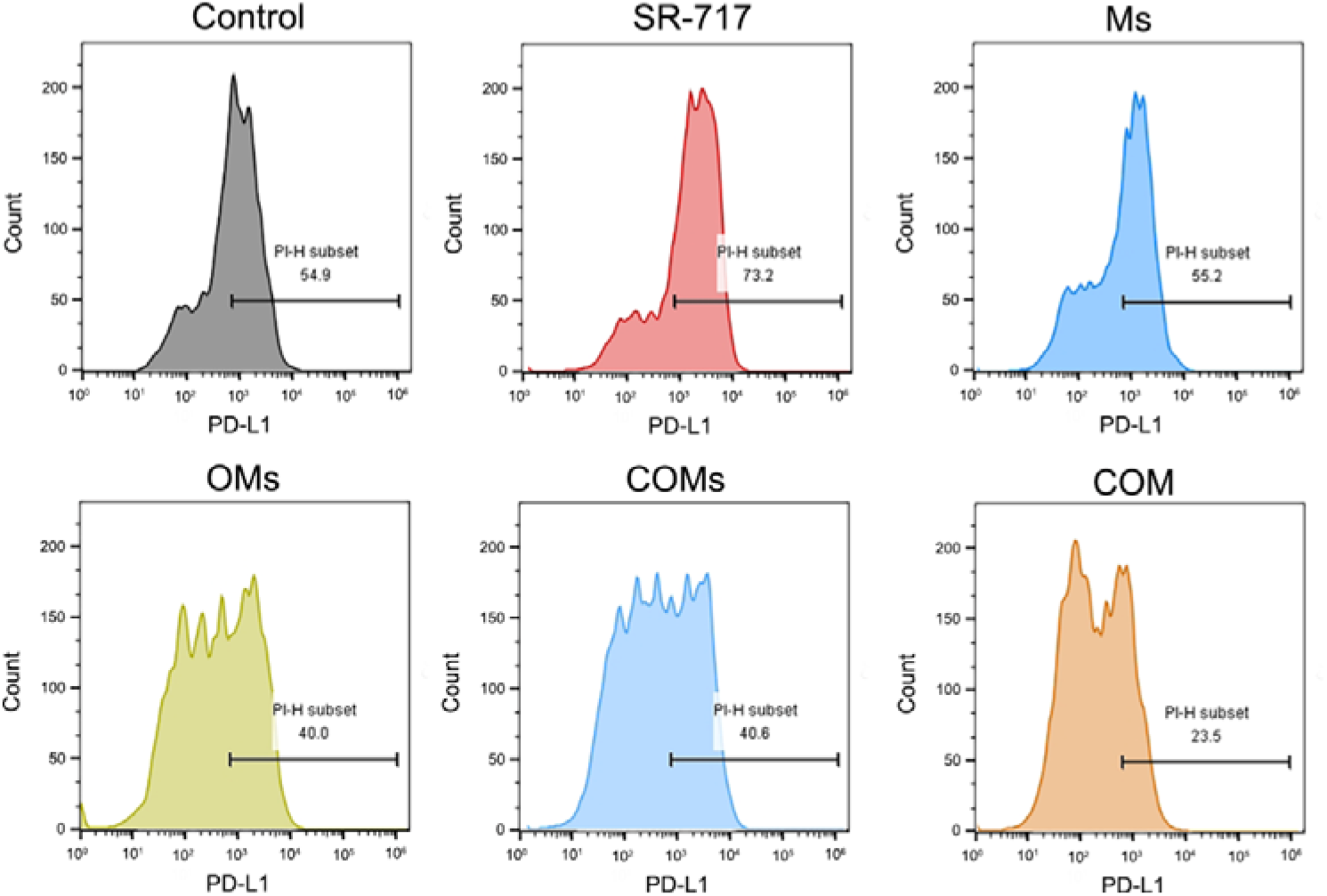
The expression of PD-L1 in B16F10 cells. B16F10 cells were treated with SR-717 (20 μM), MS, OMs, COMs or COM for 18h, and the expression of PD-L1 in tumor cells was detected by flow-cytometry. n=5.

**Figure S7.**
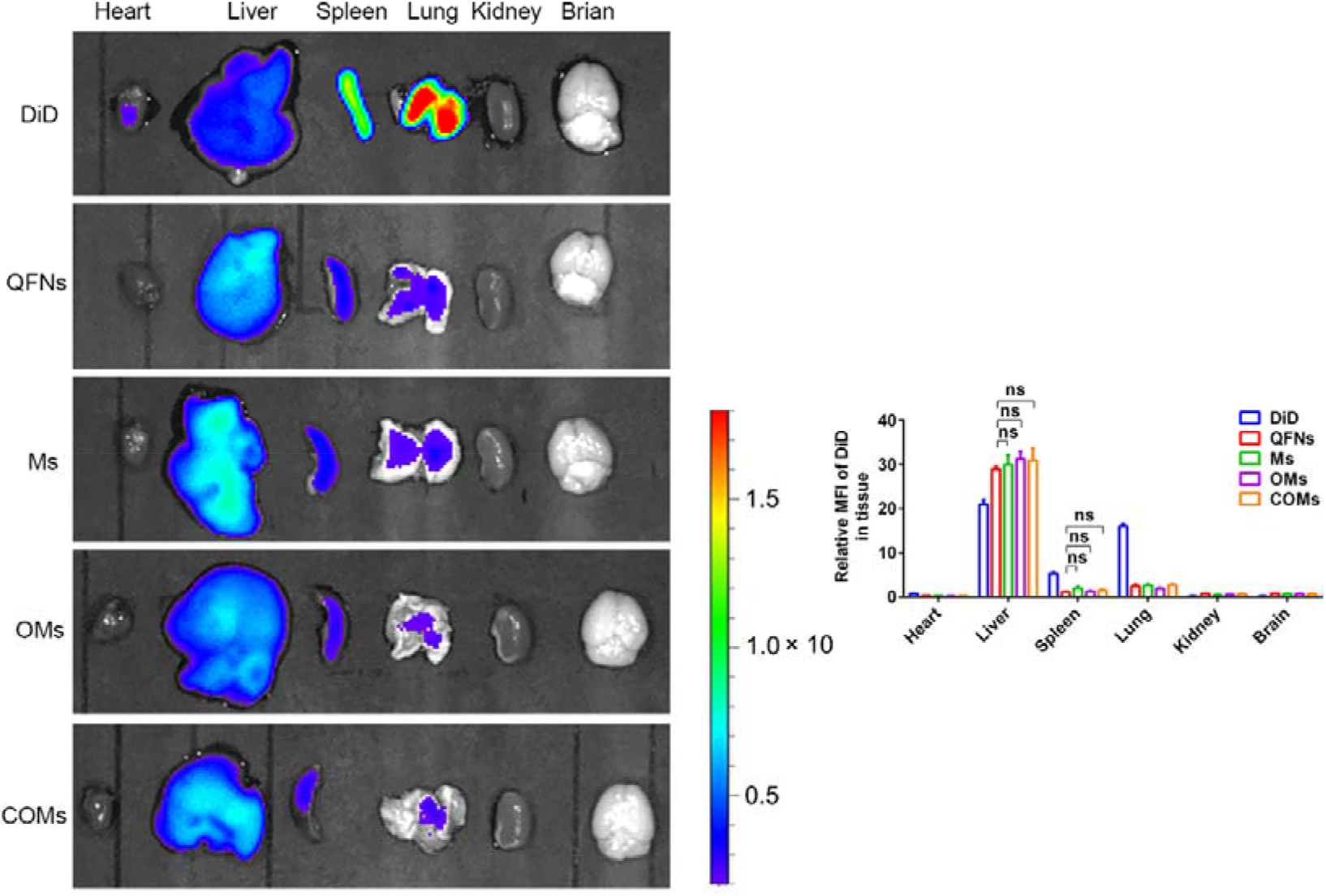
Fluorescence images of major organs after treated with free DiD, QFNs, Ms, OMs and COMs at 24 h post injection (iv.). The semi-quantitative analysis is shown on the right. n=3.

**Figure S8.**
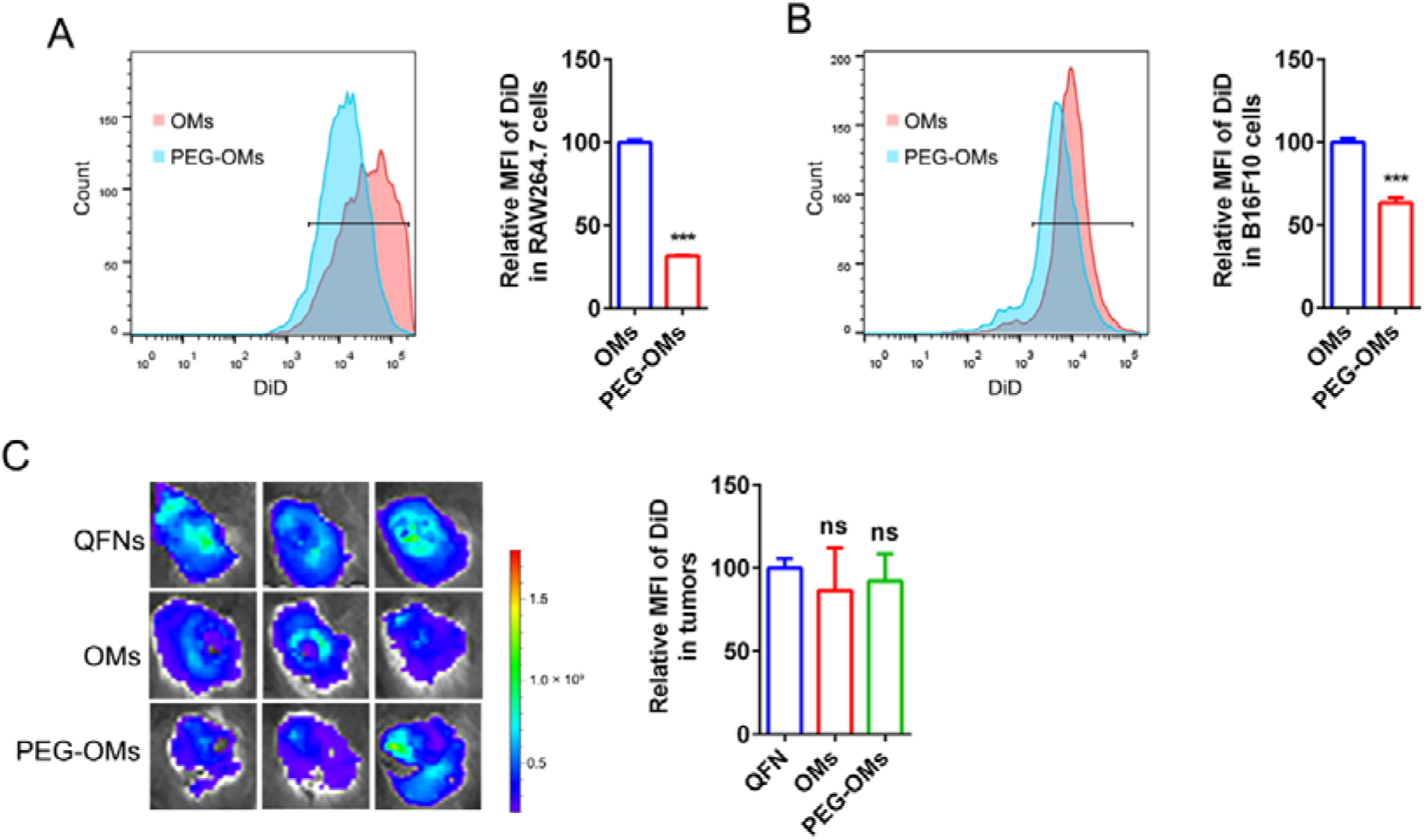
The tumor targeting ability of PEG-OMs in vivo. (A)(B) The uptake of PEG-OMs by RAW264.7 cells and B16F10 cells detected by flow-cytometry. The semi-quantitative analysis is shown on the right. n=5. (C) The fluorescence images of B16F10 tumor in mice treated with QFNs, OMs and PEG-OMs at 12 h post injection (iv.), and the semi-quantitative analysis is shown on the right. n=3.

**Figure S9.**
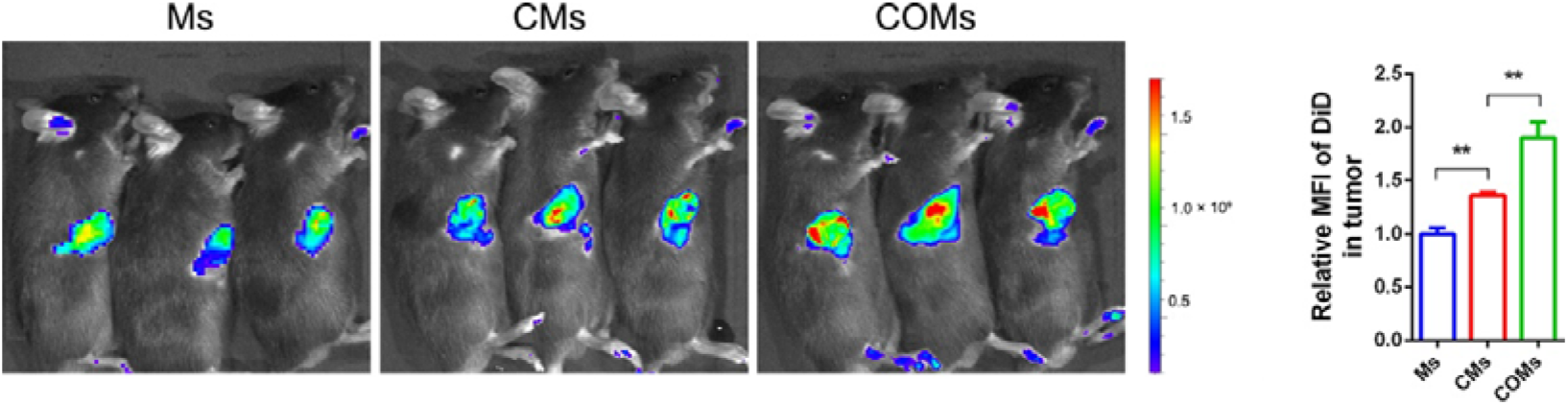
Fluorescence images of tumor in mice treated with Ms, CMs and COMs at 12 h post injection (iv.). The semi-quantitative analysis is shown on the right. n=3.

**Figure S10.**
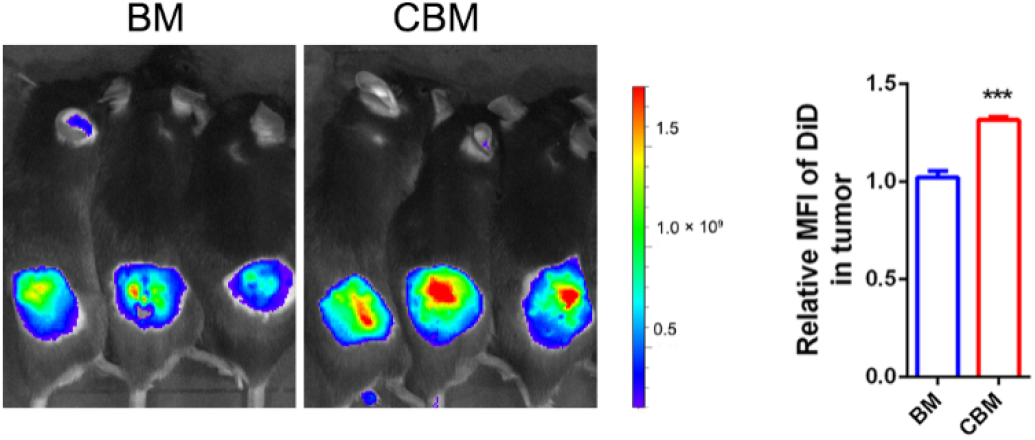
Fluorescence images of tumor in mice treated with BM and CBM at 12 h post injection (iv.). The semi-quantitative analysis is shown on the right. n=3.

**Figure S11.**
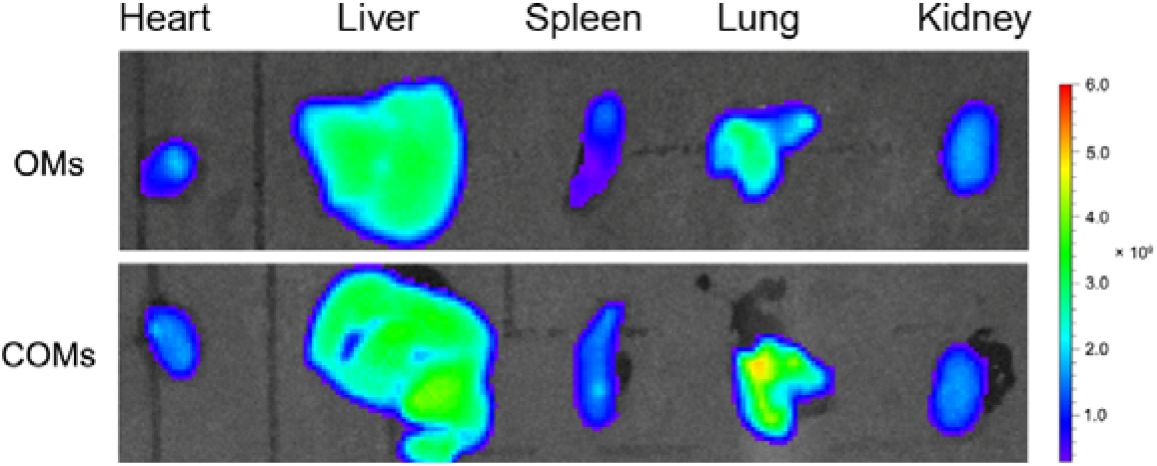
Representative fluorescence images of major organs after treated with free OMs and COMs at 0.5 h post injection (iv.). n=5.

**Figure S12.**
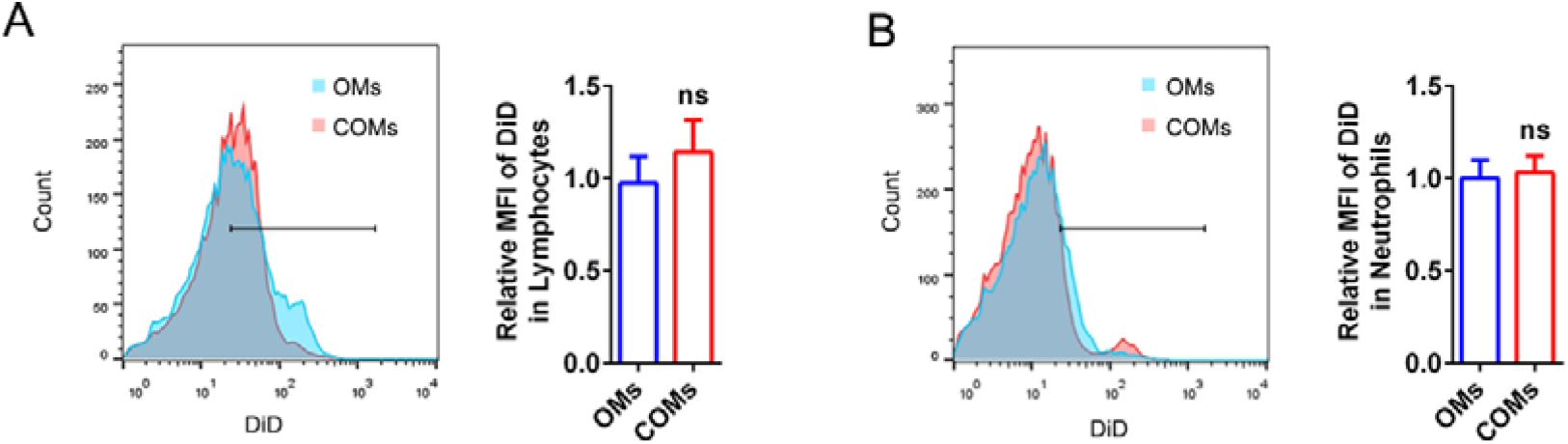
The uptake of OMs and COMs by lymphocytes and neutrophils. (A) The uptake of OMs and COMs by lymphocytes in the blood at 0.5 h post injection (iv.). and the semi-quantitative analysis is shown on the right. n=5. (B) The uptake of OMs and COMs by neutrophils in the blood at 0.5 h post injection (iv.), and the semi-quantitative analysis is shown on the right. n=5.

**Figure S13.**
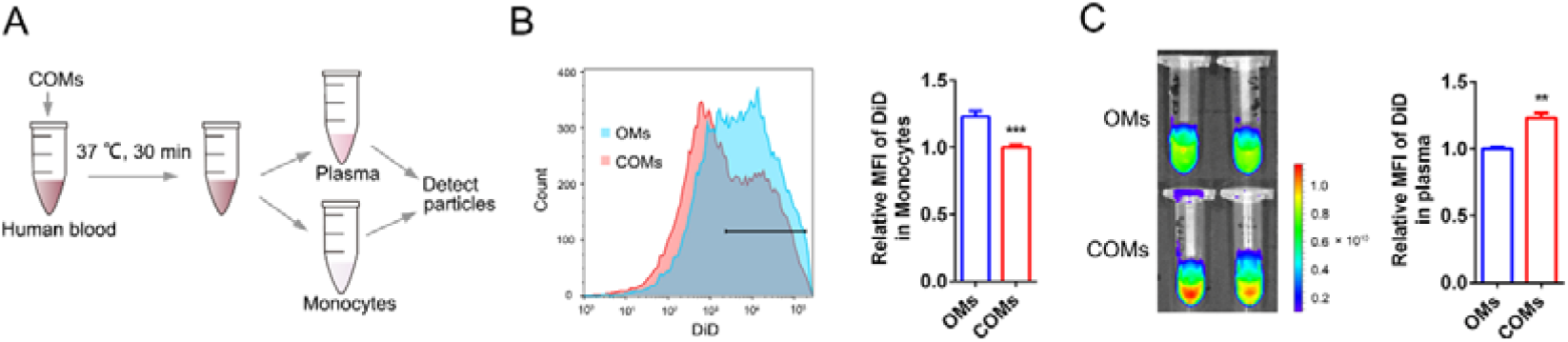
The uptake of OMs and COMs by monocytes in human blood. (A) Schematic illustration of the experiment process. (B) The uptake of OMs and COMs by monocytes in the blood at 30 min post incubation, and the semi-quantitative analysis is shown on the right. n=6. (C) Representative fluorescence images of plasma at 30 min post incubation, and the semi-quantitative analysis is shown on the right. n=6.

**Figure S14.**
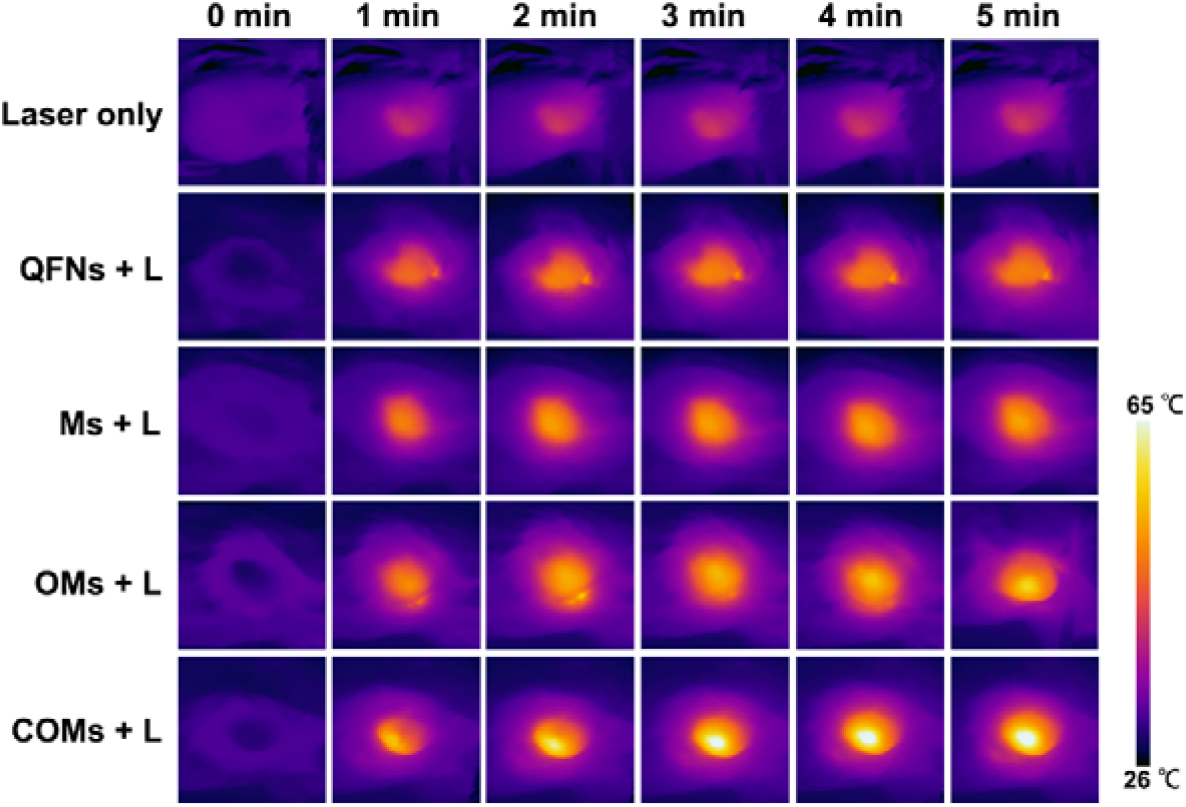
The infrared thermal images of mice. Mice bearing tumors were injected with QFNs, Ms, OMs or COMs, and mice were irradiated for 5 min with a near-infrared laser (808 nm, 1.0 W/cm^2^) at 12 h post injection (iv.). n=3.

**Figure S15.**
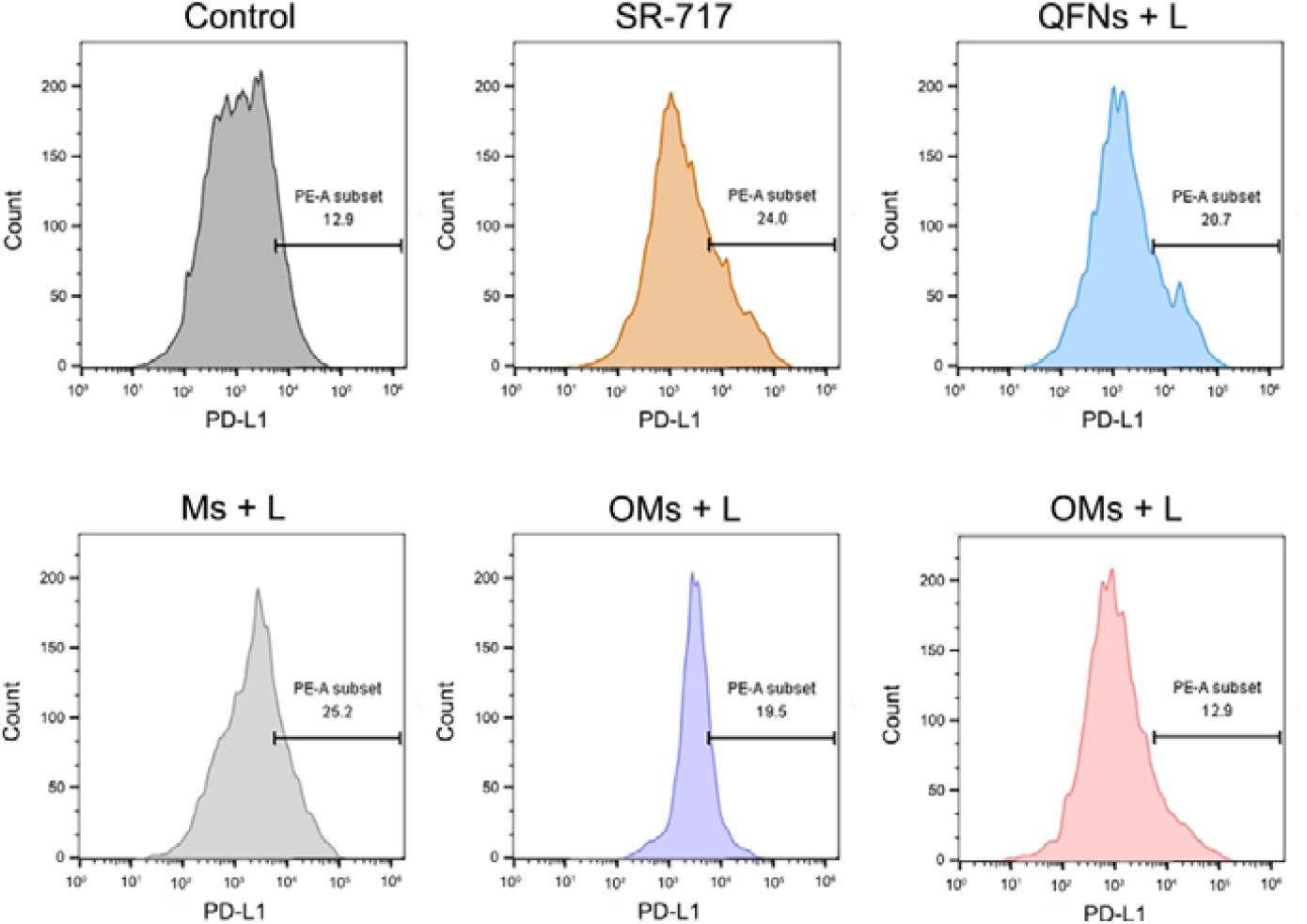
The expression of PD-L1 in B16F10 tumors at 7 days post laser irradiation. n=5.

**Figure S15.**
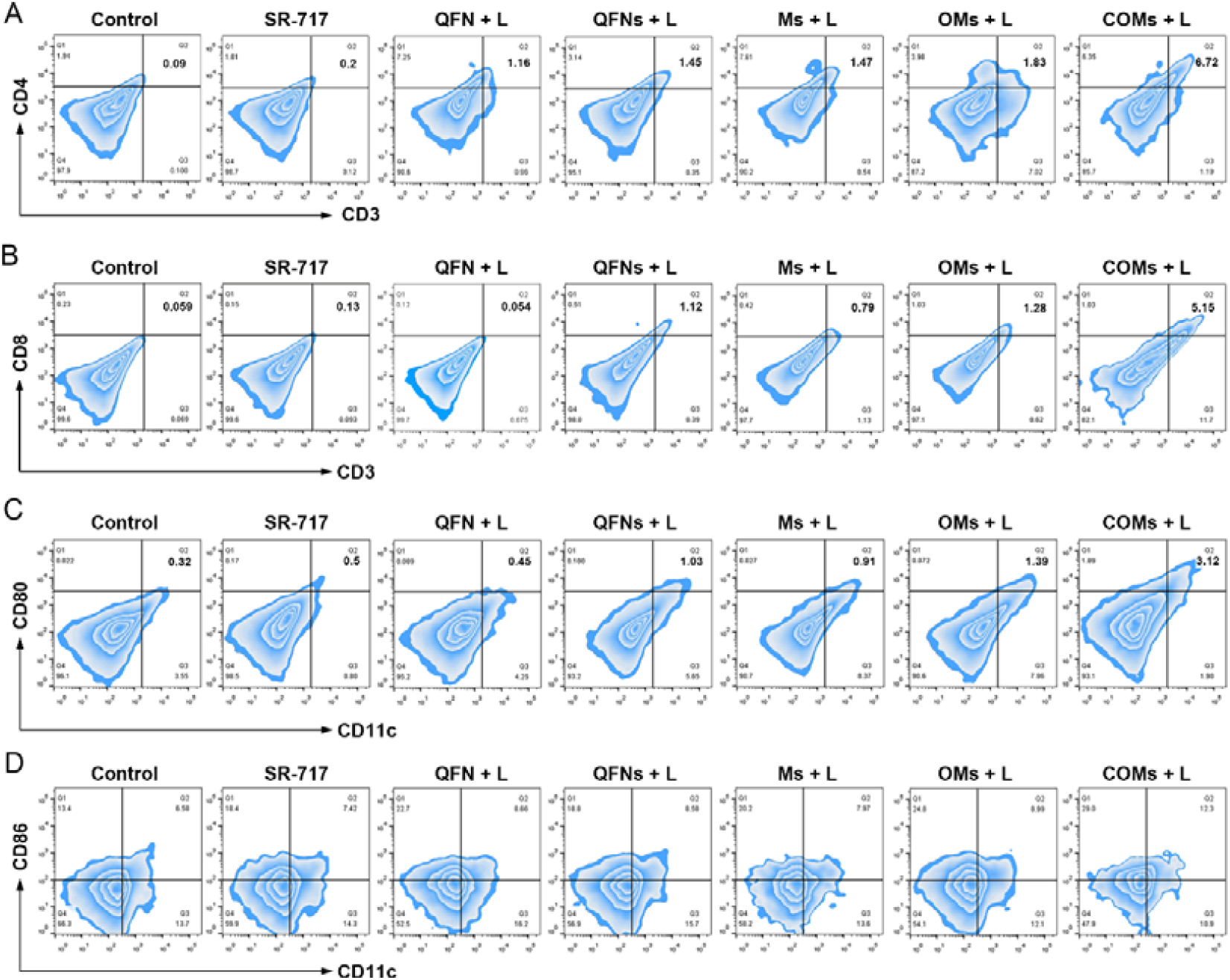
In vivo antitumor immunity activation after different treatments. (A)(B) Flow-cytometric analysis of CD4^+^T cells and CD8^+^ T cells in tumors at 7 days post laser irradiation. n=5. (C) Flow-cytometric analysis of M1 macrophages in tumors at 7 days post laser irradiation. n=5. (D) Flow-cytometric analysis of the maturation of DCs in lymph node at 7 days post laser irradiation. n=5.

**Figure S16.**
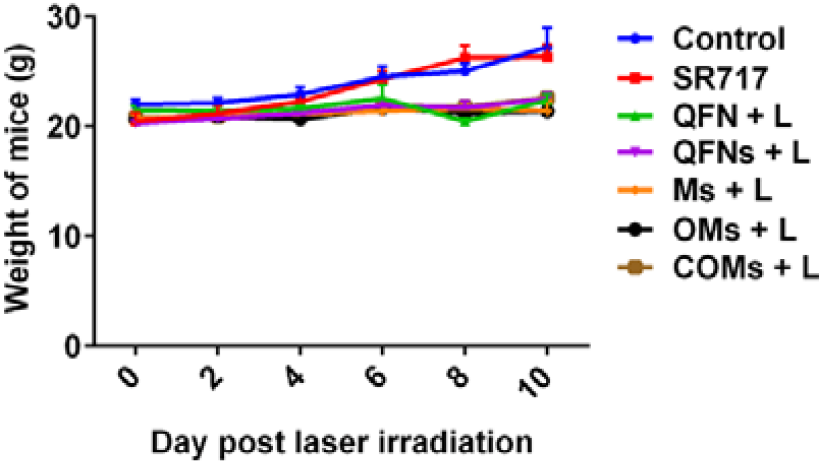
Body weight of mice after different treatments. n=5.

**Figure S17.**
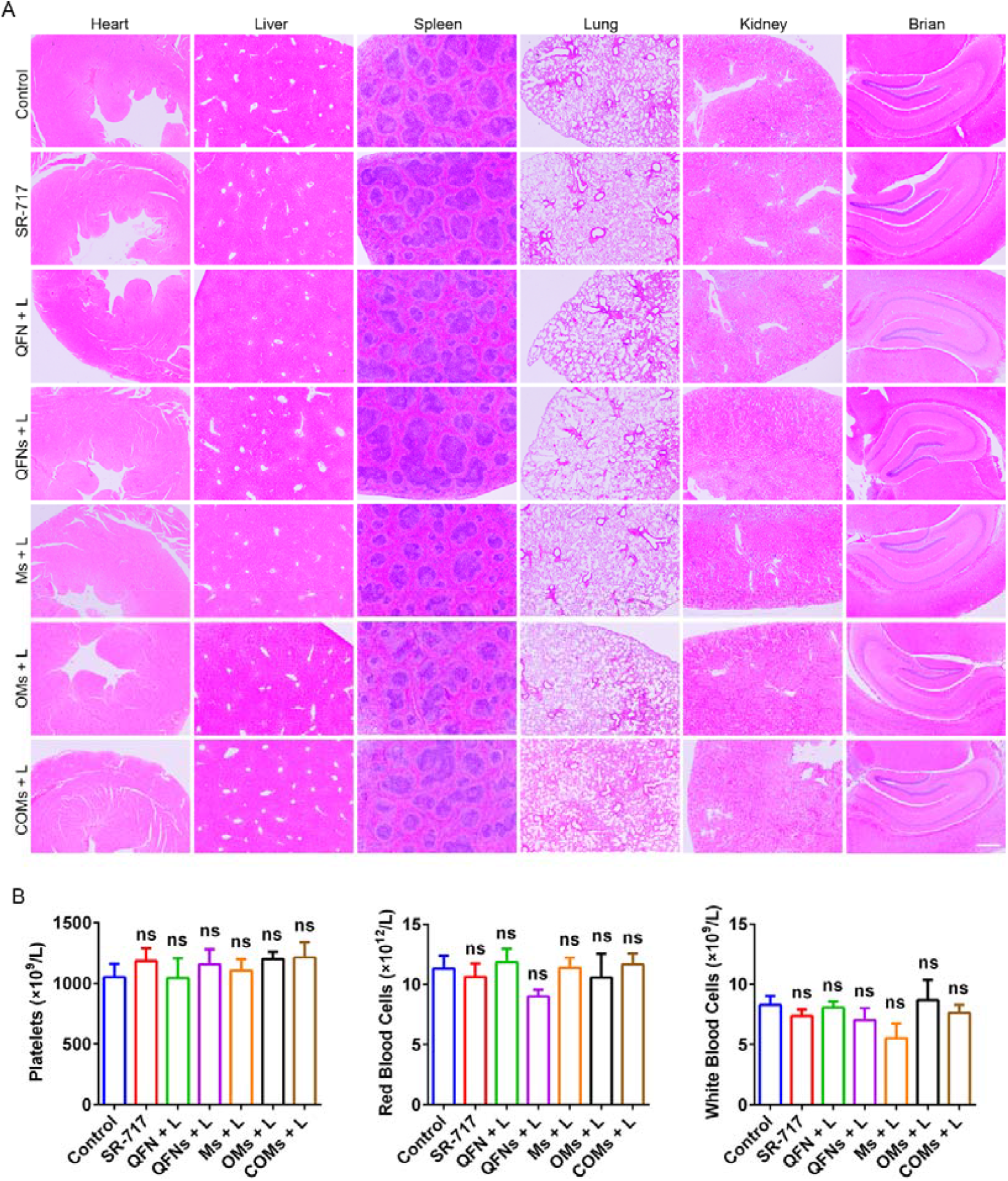
Biosafety of COMs in mice. (A) H&E staining of major organs after different treatments at 7 days post laser irradiation. n=3. Scale bar, 100 μm. (B) Blood routine examination of mice at 7 days post laser irradiation. n=5.

